# Emergent properties in complex synthetic bacterial promoters

**DOI:** 10.1101/152934

**Authors:** Lummy Maria Oliveira Monteiro, Letícia Magalhães Arruda, Rafael Silva-Rocha

## Abstract

Regulation of gene expression in bacteria results from the interplay between transcriptional factors (TFs) at target promoters, and how the arrangement of binding sites determines the regulatory logic of promoters is not well known. Here, we generated and fully characterized a library of synthetic complex promoters for the global regulators, CRP and IHF, in *Escherichia coli*, formed by a weak -35/-10 consensus sequence preceded by four combinatorial binding sites for these TFs. We found that while *cis*-elements for CRP preferentially activate promoters when located immediately upstream of the promoter consensus, binding sites for IHF mainly function as “UP” elements and stimulate transcription in several different architectures in the absence of this protein. However, the combination of CRP- and IHF-binding sites resulted in emergent properties in these complex promoters, where the activity of combinatorial promoters cannot be predicted from the individual behavior of its components. Taken together, the results presented here add to the information on architecture-logic of complex promoters in bacteria.

## Introduction

The experience of the last decade has greatly increased our knowledge of how cells coordinate gene expression in response to changing environmental and physiological conditions. Since the seminal description of the first gene regulatory mechanism by Jacob and Monod in the 60’s, thousands of molecular studies have described the different mechanisms by which transcriptional factors (TFs) coordinate gene expression in bacteria. In particular, the model organism *Escherichia coli* has been used for decades to investigate the different ways in which TFs activate or repress gene expression and a number of mechanisms have been elucidated in this and other bacteria (Browning & Busby, 2004b; Collado-Vides et al, 1991; De Lorenzo et al, 1988; Little et al, 1980; Liu & Matsumura, 1994; Miyada et al, 1984; Prigent-Combaret et al, 2001; Vicente et al, 1999). With an increase in our knowledge of these mechanisms, it was soon evident that bacterial promoters are usually regulated by several TFs that bind to specific *cis*-regulatory elements located in close proximity to the promoter site and interact with one another in different ways. In this sense, the existence of synergy or competition between TFs for binding sites in the DNA will ultimately determine the level and timing of expression for each particular gene depending on the combination of specific molecular signals available to the bacteria (Buchler et al, 2003; Hermsen et al, 2010). Additionally, compilation of the regulatory interactions known for *E. coli* resulted in the classification of TFs as global and local regulators, where the first group is composed of TFs capable of controlling a large number of target genes, whereas the second group has a more limited regulatory scope (Martinez-Antonio & Collado-Vides, 2003; Pérez-Rueda & Collado-Vides, 2000). This analysis also showed that some environmental and physiological signals that control global regulators are higher in the regulatory hierarchy since their presence will lead to major regulatory effects in the organisms compared to the presence of signals for local regulators. For instance, the CRP global regulator controls the expression of a larger number of genes in *E. coli* in response to changes in cAMP levels (which in turn is modulated by glucose (Inada et al, 1996; Schmitz, 1981)). In another case, the nucleoid associated protein IHF has an important role in DNA organization in response to bacterial growth and can modulate the expression of a number of genes (Azam & Ishihama, 1999; Biek & Cohen, 1989). Furthermore, many global regulators are known to co-occur frequently at target promoters (Collado-Vides et al, 1991; Martinez-Antonio & Collado-Vides, 2003), and this co-occurrence could indicate the existence of some interaction mechanisms between these pairs of regulators (Browning & Busby, 2004a; Guazzaroni & Silva-Rocha, 2014; Lee et al, 2012).

With the advent of synthetic biology, using the current knowledge on gene regulatory mechanisms in bacteria to reprogram these organisms for novel applications has been of special interest (Benner & Sismour, 2005; Nandagopal & Elowitz, 2011; Voigt, 2006). In order to accomplish this task, many studies have addressed the modification of native promoters to construct synthetic regulatory systems with enhanced and/or modified performance (Brophy & Voigt, 2014; Gardner et al, 2000; Guet et al, 2002; Rhodius et al, 2013). Moreover, some initial studies have focused on the shuffling of *cis*-regulatory elements to re-construct complex promoters in bacteria (Cox et al, 2007; Isalan et al, 2008; Kinkhabwala & Guet, 2008; Murphy et al, 2007); this approach could not only provide novel regulatory systems but also reveal some of the hidden roles regarding the interaction of multiple TFs in target promoters. Though these approaches have resulted in significant progress such as the knowledge that promoter arrangement indeed determines the final regulatory logic of systems, these studies have mainly used local regulators and it is not yet known whether global regulators would follow the same rules.

Moreover, the standard model for gene regulation in bacteria states that we could anticipate the regulatory behavior of complex promoters by analyzing the individual contributions of each TF and its respective *cis*-elements, as evidenced by the widely used mathematical frameworks available to model gene regulation (Bintu et al, 2005a; Bintu et al, 2005b). However, in an alternative scenario, the combination of several *cis*-regulatory elements for specific TFs (mainly for global regulators that naturally act together) could result in promoters with emergent properties, where the final response of the system would not be anticipated based on known individual contributions. This hypothesis is also motivated by the fact that many biological systems have been shown to display emergent properties (Bhalla & Iyengar, 1999).

In order to get insights into the regulatory mechanisms of combinatorial bacterial promoters, we investigated here the relationship between promoter architecture and gene expression regulation. For this, we constructed and characterized a library of synthetic promoters containing an array of *cis*-elements for CRP and IHF, two global regulators of *E. coli*, using a GFP reporter assay. Our data clearly indicated that though CRP and IHF have very different regulatory effects, many binding site combinations for both TFs resulted in novel regulatory activities that were not anticipated by the analysis of individual elements when their sites were placed in different positions and arrangements. These results demonstrate the existence of emergent properties in complex synthetic promoters in *E. coli*, which could be extrapolated to naturally occurring regulatory systems and would significantly impact the engineering of synthetic biological circuits in bacteria.

## Results

### CRP strongly activates synthetic promoters with cis-elements immediately upstream of a core promoter

In order to investigate the architecture-logic relationship in synthetic bacterial promoters, we employed a minimal design as presented in Figure 1. First, we designed a promoter composed of a weak core element (comprising the -35 and -10 boxes of *Plac*) preceded by 20 bp sequences that could be occupied by cis-elements for the target TFs (**Fig. 1A**). In this design, the *cis*-elements can be centered at regions -61, -81, -101, and -121 related to the transcriptional start site (TSS) of the promoter. By fixing these positions, we would expect the effect of TF to be stimulatory at the resultant promoter, based on previous systematic inspections on the effect of *cis*-element localization on gene regulation (Collado-Vides et al, 1991; Tebbutt et al, 2002). For each of the four potential positions, we designed double stranded DNA oligonucleotides with a consensus sequence for CRP, IHF, and a control sequence (called “Neg”, **Fig. 1B**) that does not display any stimulatory effect *in vivo* (Guazzaroni & Silva-Rocha, 2014). In this sense, each double stranded DNA fragment has 3’ overhang elements with four nucleotides that specify each position where the fragment can be ligated (**Fig. 1C**). In this manner, we used a portfolio of 12 different fragments that could be used to construct up to 81 (3^4^) different combinatorial promoters. Using this setup, we constructed a library of synthetic promoters by ligating these *cis*-elements and the core promoter into a GFP reporter vector, allowing the measurement of promoter activities *in vivo* to determine the effect of promoter architecture in the final output of promoters (exemplified by the different clones represented in **Fig.1D**).

**Figure 1.**
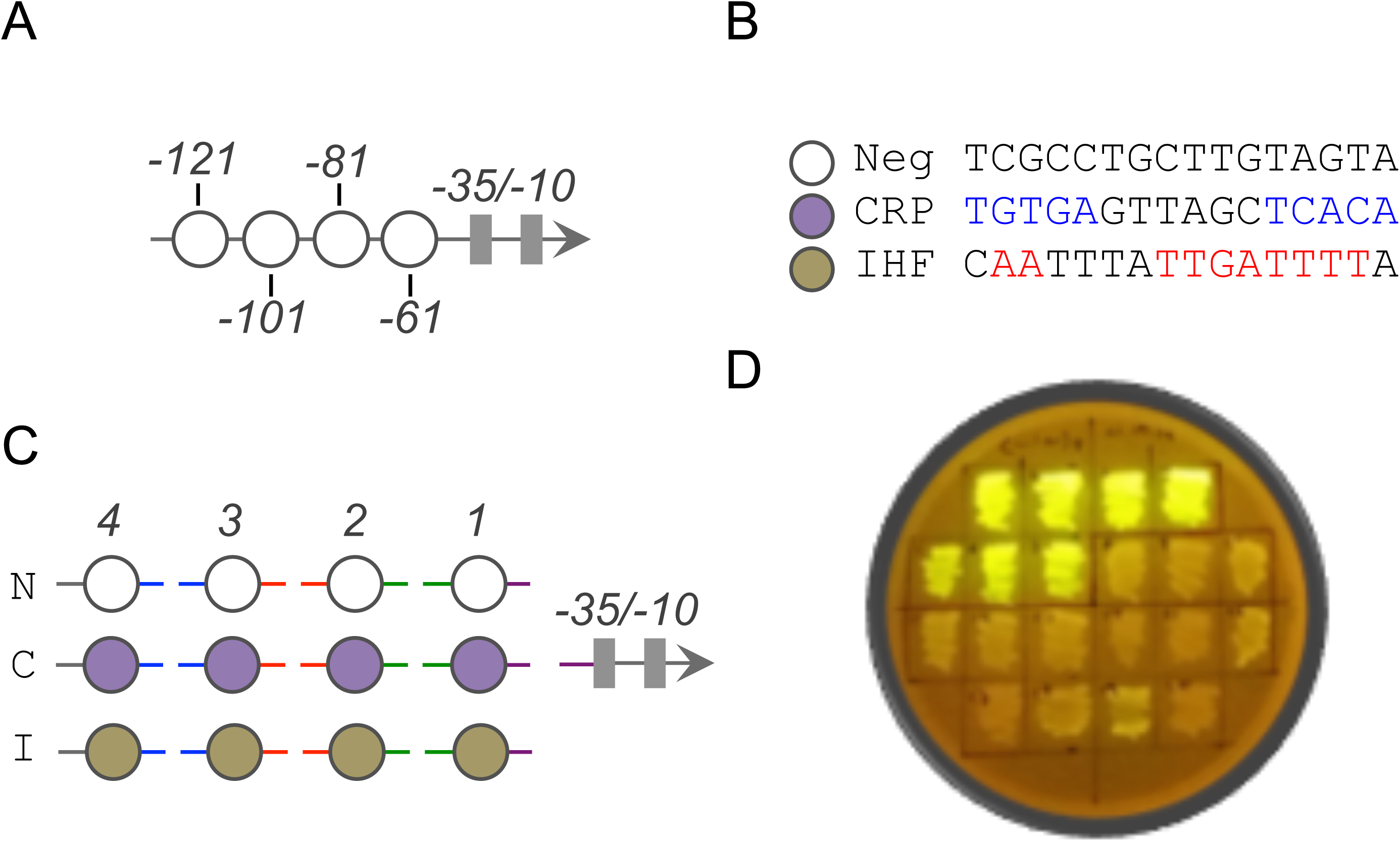
Construction of the complex promoter library. A) Schematic representation of the promoter library, showing the positions -121, -101, -81, and -61 (white circles) at which *cis*-elements were inserted. The -35 and - 10 boxes (grey rectangles) correspond to the core promoter. B) Nucleotide sequences for Neutral (N), CRP (C), IHF (I) *cis*-elements. C) Simplified scaffold scheme for the minimal synthetic promoter library. Motifs positions are identified as 4, 3, 2, and 1 respective to the core promoter, and colored lines represent the cohesive sequences for DNA ligation. D) *E. coli* library transformants showing different promoter strengths.

In order to determine the effect of different arrangements of *cis*-elements for CRP, we analyzed the promoter activity of 10 synthetic promoters in the wild type strain growing in minimal media for a period of 8 hours. As shown in **Fig. 2**, clustering of promoter activities using Euclidean distance reveals the existence of two clear groups, one (marked as **I** in the figure) composed of six promoters with activities similar to the reference promoter (the one in the top with four Neg sites) and another group (marked as **II**) composed of four promoters with a high level of activity (about 80 times the level of the reference promoter). By analyzing the architecture of each promoter, it is easily notable that all members of the highly active group have a *cis*-element for CRP at position 1 (equivalent to the -61 relative to the TSS), which is in accordance to previous reports on this TF (Tebbutt et al, 2002; Ushida & Aiba, 1990). Moreover, the addition of another CRP *cis*-element at positions 2, 3, and 4 (boxes - 81, -101, and -121) only marginally affects the activity of a promoter harboring a *cis*-element at position 1. In summary, this result demonstrates the potential of our approach to investigate the effect of *cis*-element arrangement on promoter activity, as the resulting synthetic promoters reproduced the expected behavior for CRP.

**Figure 2.**
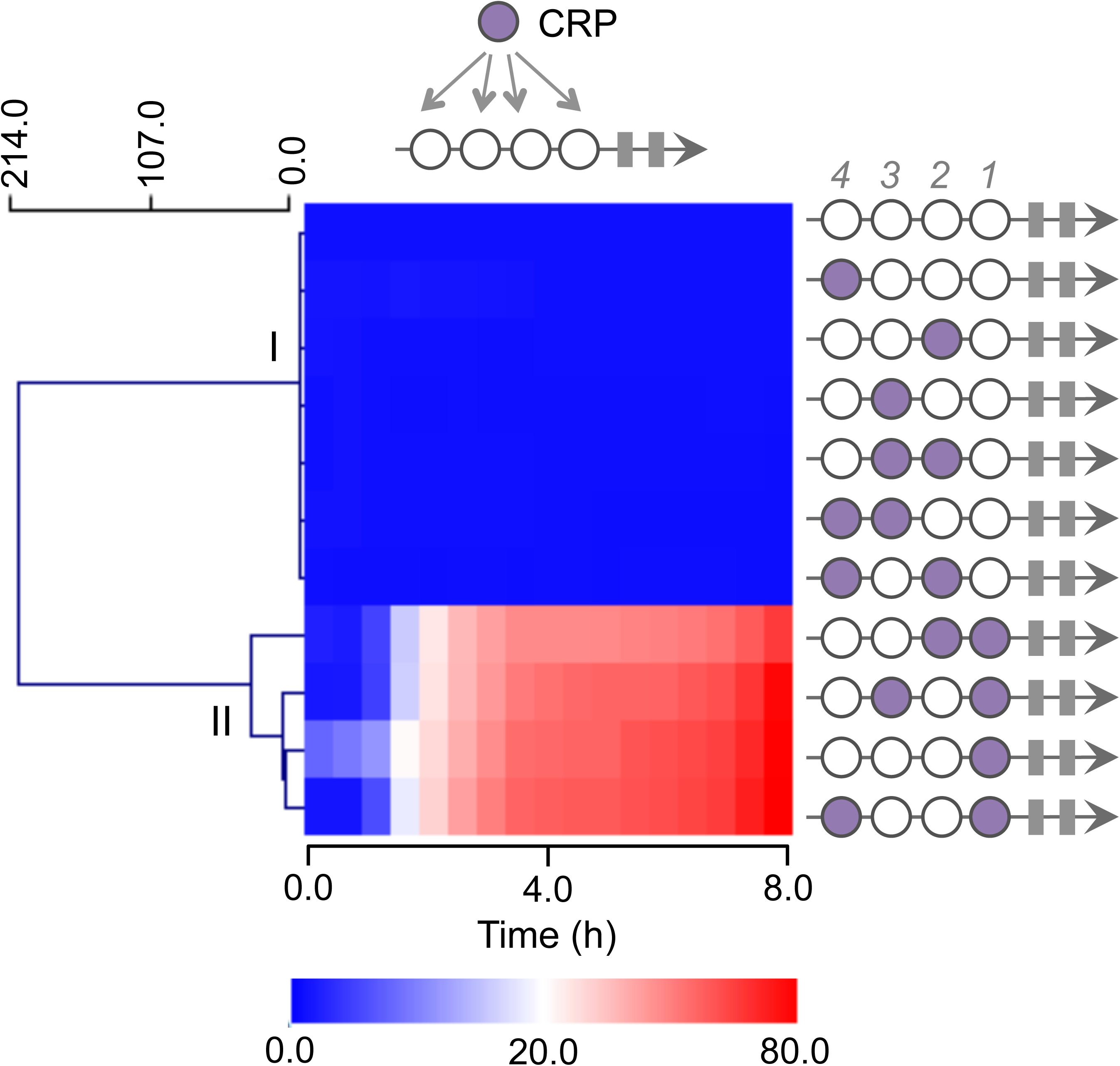
CRP motif at position 1 is fundamental for high promoter activity. A subset of 11 synthetic promoters containing shuffled CRP and Neutral *cis*-elements displaying two clear activity patterns (groups **I** and **II**). In group **I** are promoters that do not present promoter activity while group **II** includes promoters with high transcription rates. Circles in magenta represent the positions of CRP sites. Relative promoter activity was measured for 8 h, calculated based on the Neutral full promoter, and displayed on an intensity scale from 0.0 to 80.0. Plots were calculated based on the average of three independent experiments.

### cis-element for IHF enhances promoter activity in the complex promoters in the absence of this TF

In order to investigate the effect of *cis*-elements for IHF in our complex promoter design, we analyzed 11 synthetic promoters harboring combinations of IHF sites and Neg sequences (**Fig. 3**). The experiments were performed similarly as before but both in the wild type and Δ*ihf* mutant strains of *E. coli*. Promoter activity analysis allowed clustering of the data into three major groups as shown in **Fig. 3A** for the wild type strain. In this sense, group **I** was formed of two promoters with maximal activity not higher than 8 times that observed for the reference promoter, whereas group **II** (5 promoters) displayed activities comparable to those of the reference, and group **III** (4 promoters) showed intermediate activity. When the same set of promoters was assayed in Δ*ihf* mutant strains of *E. coli*, we observed two major features (**Fig. 3B**). First, a generalized increase in promoter activity was observed for groups **I** and **III**, where the former was still formed of promoters with stronger activity. Second, the composition of the groups was almost unchanged, with the exception of one promoter (with two IHF *cis*-elements at positions 4 and 3) that displayed no activity in the wild type but showed the highest activity of the group in the mutant, and another promoter (composed of four IHF binding sites) that did not gain activity in the mutant, were clustered into group **II** in the mutant. These expression profiles clearly indicate that the *cis*-elements for IHF could stimulate promoter activity mainly in the absence of this global regulator. In addition, the two groups displaying significant promoter activity in the mutant strain (groups **I** and **III**) have *cis*-elements for IHF at many locations except position 2 (equivalent to the -81 region), whereas the promoters with this position occupied, displayed very low activity regardless of occupancy at other sites (group **II**). These results suggest that *cis*-elements for IHF could operate as an RNAP transcriptional activity enhancer, probably as a UP element-like motif as described previously (Giladi et al, 1996; Rossiter et al, 2015). Thus, when IHF binds to its cognate *cis-*element, it blocks RNAP contact with the UP element-like sequence, thus preventing transcriptional stimulation of the promoter. However, the reason why the existence of a *cis*-element for IHF at position 2 renders the promoters inactive regardless of the identity of other positions is unknown and may be related to some intrinsic property of the DNA sequence itself.

**Figure 3.**
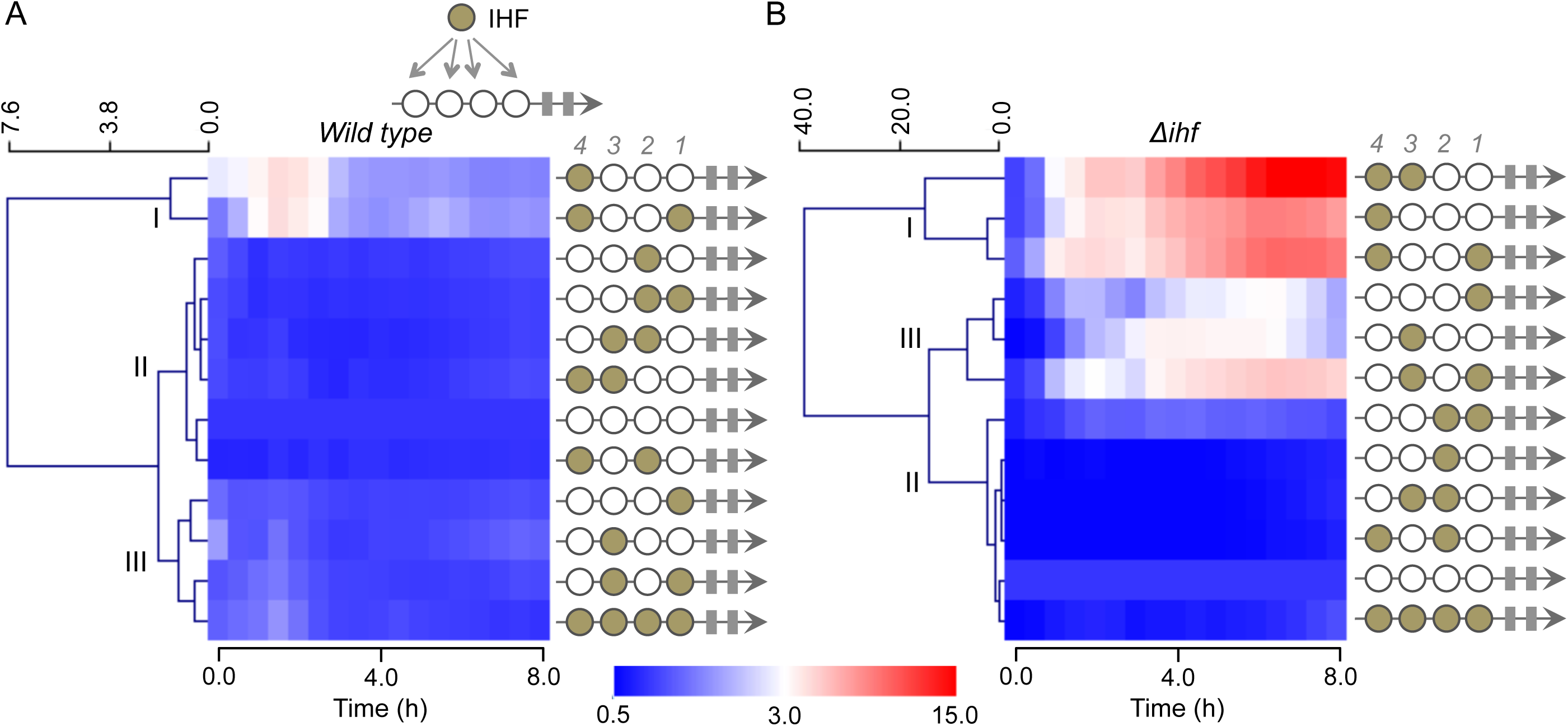
IHF motif enhanced promoter activity in *E. coli* Δ*ihf* strain. A subset of shuffled IHF and Neutral motif promoters were assayed in the wild type and Δ*ihf* mutant strains and grouped according to their relative activity. Circles in beige represent the positions of IHF sites. A) IHF *vs.* Neutral motifs assayed in the wild type strain. Synthetic promoters that showed higher promoter activities are clustered in group **I**, group **II** is formed of promoters with low activity, whereas group **III** is formed of promoters with intermediate promoter activity. B) The same set of promoters were assayed in the *E. coli* Δ*ihf* mutant strain, highlighting that in the absence of IHF transcription factor, promoter activity was generally improved for the groups **I** and **III**. Relative promoter activity was measured for 8 h, calculated based on the Neutral full promoter, and displayed on an intensity scale from 0.0 to 15.0. Plots were calculated based on the average of three independent experiments.

### Rise of emergent properties in complex promoters with CRP and IHF cis-elements

Once we determined that the CRP and IHF *cis*-elements have distinct effects on promoter activity, we wondered what would happen if the CRP and IHF binding sequences were combined, as occurs in the natural promoters of *E. coli*. In this sense, would the resulting promoter represent the sum of each contribution of the isolated *cis*-elements, or would it display a novel regulatory logic? To address these questions, we used as the start point, two architectures containing *cis*-elements for IHF that displayed activity both in the wild type and Δ*ihf* mutant strains as represented by the members of group **I** in **Fig. 3A**. These two promoters possess either one position (position 4) or two (positions 4 and 1) occupied by the IHF *cis*-element. Using these two basic architectures, we introduced *cis*-elements for CRP at either position 3, 2, or both, and assayed the resulting promoter activity (**Fig. 4**). In this dataset, we did not test position 1 since CRP at this position has a strong stimulatory effect regardless of other upstream elements (as already presented in **Fig. 2**). Notably, when positions 3 and 2 where occupied by CRP *cis*-elements (either in isolation or simultaneously), no significant promoter activity was detected (**Fig. 2**). When we assayed combinatorial promoters based on single IHF *cis*-elements in the wild type strain, we observed that the introduction of single or double *cis*-elements for CRP resulted in complete abolishment of promoter activity. When these promoters were evaluated in the presence of 0.4% glucose (which in our tested condition, is sufficient to block CRP activity (Gerardo Ruiz Amores & Rafael, 2015)), we did not recover the original promoter activity, suggesting that this effect was not dependent on CRP binding but rather on the combination of the DNA elements itself. When we performed a similar analysis on the variants of the promoters harboring the two IHF *cis*-elements (at positions 4 and 1), we observed a remarkably different behavior. When single CRP *cis*-elements were placed individually at positions 3 or 2, we observed the same promoter blocking effect as described previously, and this effect was not alleviated in the presence of glucose (**Fig. 4**). However, when both positions, 3 and 2, where occupied by *cis*-elements for CRP resulting in a synthetic promoter with two IHF sites flanking two CRP sites, we observed a strong increase in promoter activity compared to that of the original promoter. Interestingly, the addition of glucose resulted in complete abolishment of promoter activity, indicating that the strong enhancement of promoter activity was indeed dependent on CRP activity. These data strongly support the notion that the combination of *cis*-elements for global regulators such as CRP and IHF leads to the appearance of emergent properties, since the final regulatory behavior of the complex promoter does not represent the sum of the behavior of the original architectures (i.e., the promoter harboring two IHF sites at positions 4 and 1 and the promoter harboring two CRP sites at positions 3 and 2).

**Figure 4.**
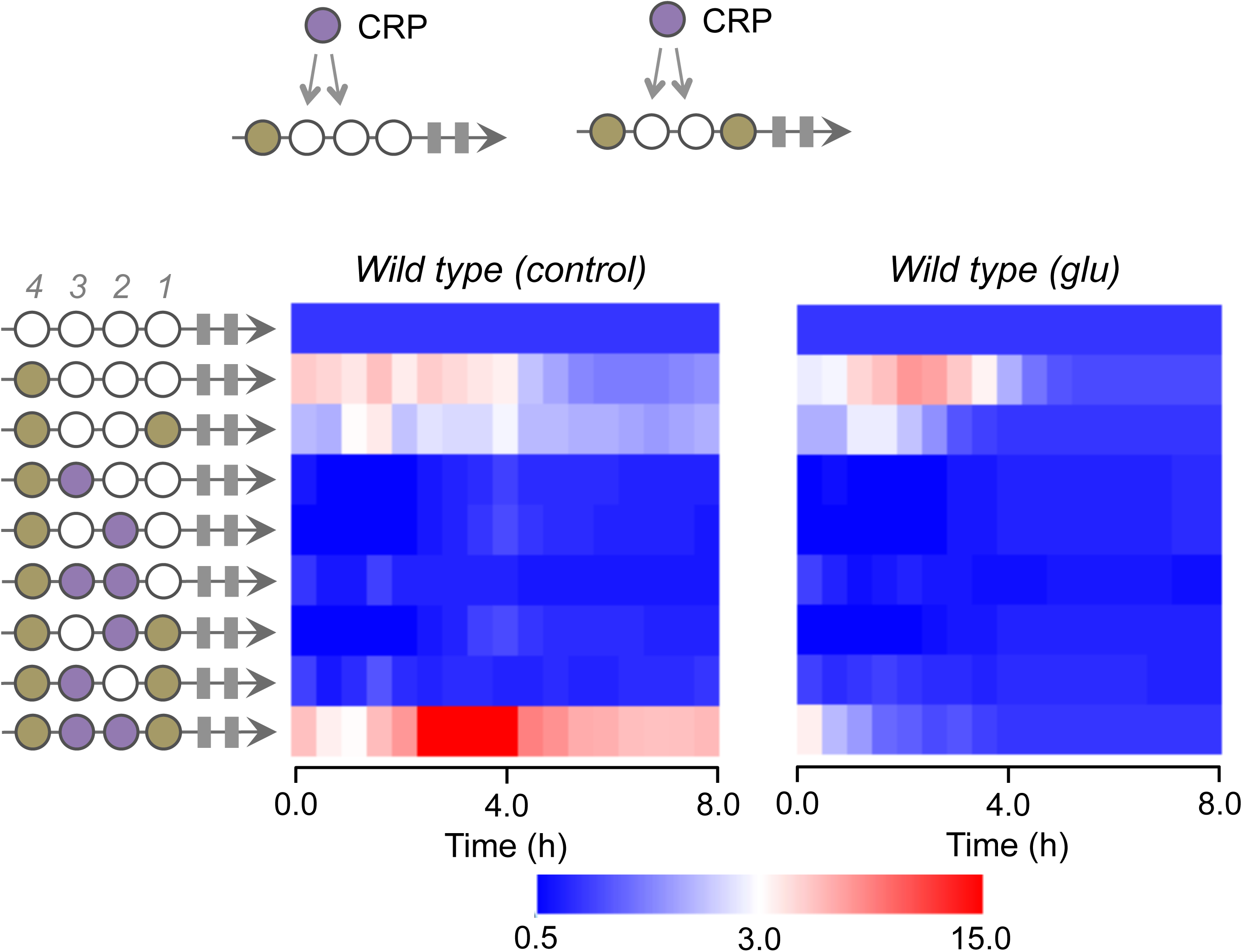
Emergence in combinatorial promoters based on CRP and IHF *cis*-elements. Complex promoters were constructed based on two promoters shown in Fig. 3 and were assayed in the *E. coli* wild type strain, in the absence (left) or presence (right) of 0.4% glucose. The promoters were constructed by fixing the IHF *cis*-element at position 4 and at positions 4 and 1 and by shuffling the CRP motifs at positions 2 and 3. Relative promoter activity was measured for 8 h, calculated based on the Neutral full promoter, and displayed on an intensity scale from 0.0 to 15.0. Plots were calculated based on the average of three independent experiments.

In order to further evaluate the promoter architecture effect, we expanded the number of architectures assayed and performed experiments in the absence of IHF and by modulating CRP activity (i.e., in the presence or absence of glucose). For a better presentation of the results, the experiment was divided into three subgroups. **Fig. 5A** shows constructs that have one IHF *cis*-element fixed at position 4 (-121 region) and different combinations of CRP *cis*-element at positions 2 and 3 (-81 and -101 regions). As shown in the figure, a promoter containing a single IHF site at position 4 displays strong activity in the Δ*ihf* mutant strain that was insensitive to glucose presence. Moreover, addition of single or double CRP cis-elements at positions 3 and 2 completely abolishes promoter activity, and this could not be reverted by the addition of glucose to the media. These results agree with the previous analysis and indicate that addition of CRP *cis*-elements blocks the activity of the original promoter independently of CRP activity. Next, we investigated the effect of the presence of CRP binding sites at different positions in promoters with a single IHF *cis*-element fixed at position 1 (**Fig. 5B**). In this condition, though the initial promoter displayed detectable activity with a fold-change about three times that of the reference promoter, addition of a single CRP *cis*-element immediately upstream of the IHF site (position 2) completely abolished the promoter activity. Interestingly, moving the CRP site far from the IHF site (for instance, from position 2 to position 3) generates a marginally detectable activity, whereas placing the site in the farthest position (i.e., at position 4) generates a combinatorial promoter with activity similar to that of the original harboring a single IHF site at position 1 (**Fig. 5B**). These results indicate a position dependent effect of CRP *cis*-elements, which was not related to CRP activity since the addition of glucose to the media resulted in very similar expression profiles. This notion is also supported by the fact that addition of two CRP binding sites at positions 4 and 3 resulted in similar activity to that where only position 3 was occupied by a CRP *cis*-element. Additionally, introduction of two CRP binding sites at positions 3 and 2 generated completely different behavior, resulting in a promoter with strong activity that was completely dependent on CRP (as addition of glucose substantially decreased its activity, **Fig. 5B**). These results indicate that the emergence of strong CRP-dependent activity requires two tandem CRP binding sites (at positions 3 and 2) followed by a single IHF *cis*-element (at position 1). Finally, when the CRP binding sites were combined with IHF *cis*-elements fixed at position 4 and 1 (**Fig. 5C**), we observed the same behavior presented in **Fig. 5B**, since addition of one site at position 3 or 2 strongly impairs promoter activity whereas two tandem sites generate a CRP-dependent promoter with activity stronger than that of the parental architecture.

**Figure 5.**
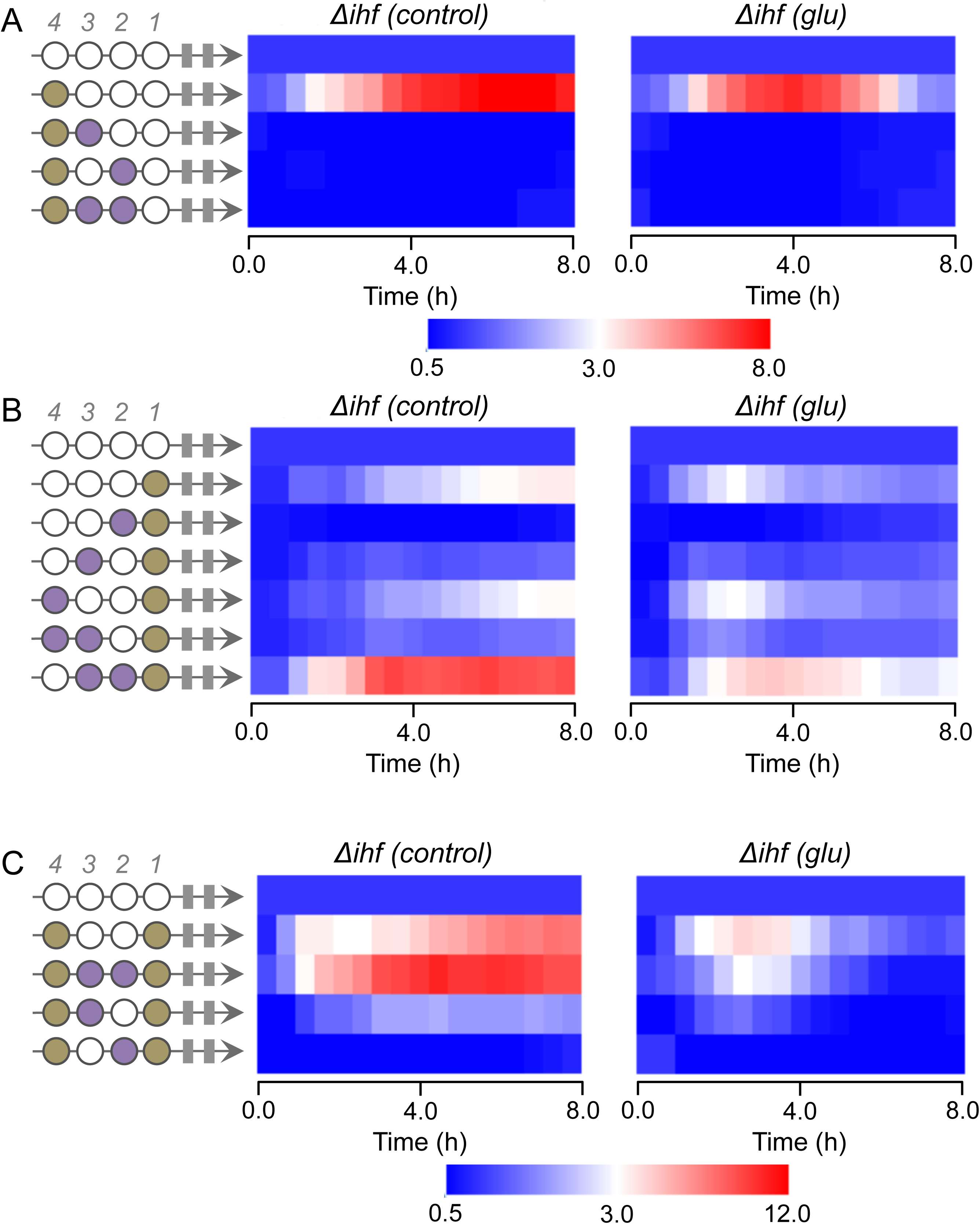
Systematic investigation of complex promoters for CRP and IHF in the *E. coli* Δ*ihf* strain. All experiments were performed in the absence (left) or presence (right) of 0.4% glucose. A) Synthetic promoters with a single IHF site fixed at position 4 and varying CRP sites at positions 3 and 2. B) Synthetic promoters with a single IHF site fixed at position 1 and varying CRP sites at positions 4, 3, and 2. C) Synthetic promoters with two IHF sites fixed at positions 4 and 1 and varying CRP sites at positions 3 and 2. Relative promoter activity was measured for 8 h, calculated based on the Neutral full promoter, and displayed on an intensity scale from 0.0 to 8.0 (at A) or 0.0 to 12.0 (at B and C). Plots were calculated based on the average of three independent experiments.

### IHF cis-elements generates fine-tuning for CRP activated promoters

In the previous sections, we demonstrated that additional CRP *cis*-elements could influence the regulatory behavior of a promoter harboring IHF sites. Since CRP binding sites could strongly influence these promoters, we investigated how the addition of IHF could modulate promoters containing CRP *cis*-elements at position 1, which were previously demonstrated to generate strong CRP dependent activation (**Fig. 2**). For this, we sampled several combinatorial promoters where additional IHF and CRP sites were mixed upstream of a CRP *cis*-element located at position 1. As shown in **Fig. 6A**, all promoters displayed strong activity in the wild type strain of *E. coli* and this activity was severely impaired when glucose was added to the growth media. Additionally, certain degree of heterogeneity can be observed in promoter activities indicating that the additional sites contributed to the final activity observed. However, when the same experiments were performed in a Δ*ihf* mutant strain of *E. coli*, we observed that the level of heterogeneity was strongly reduced both in the active and repressed conditions (**Fig. 6B**). Taken together, these data strongly indicate that additional IHF sites could produce a fine-tuning effect that modulates the final activity of the constructed combinatorial promoters.

**Figure 6.**
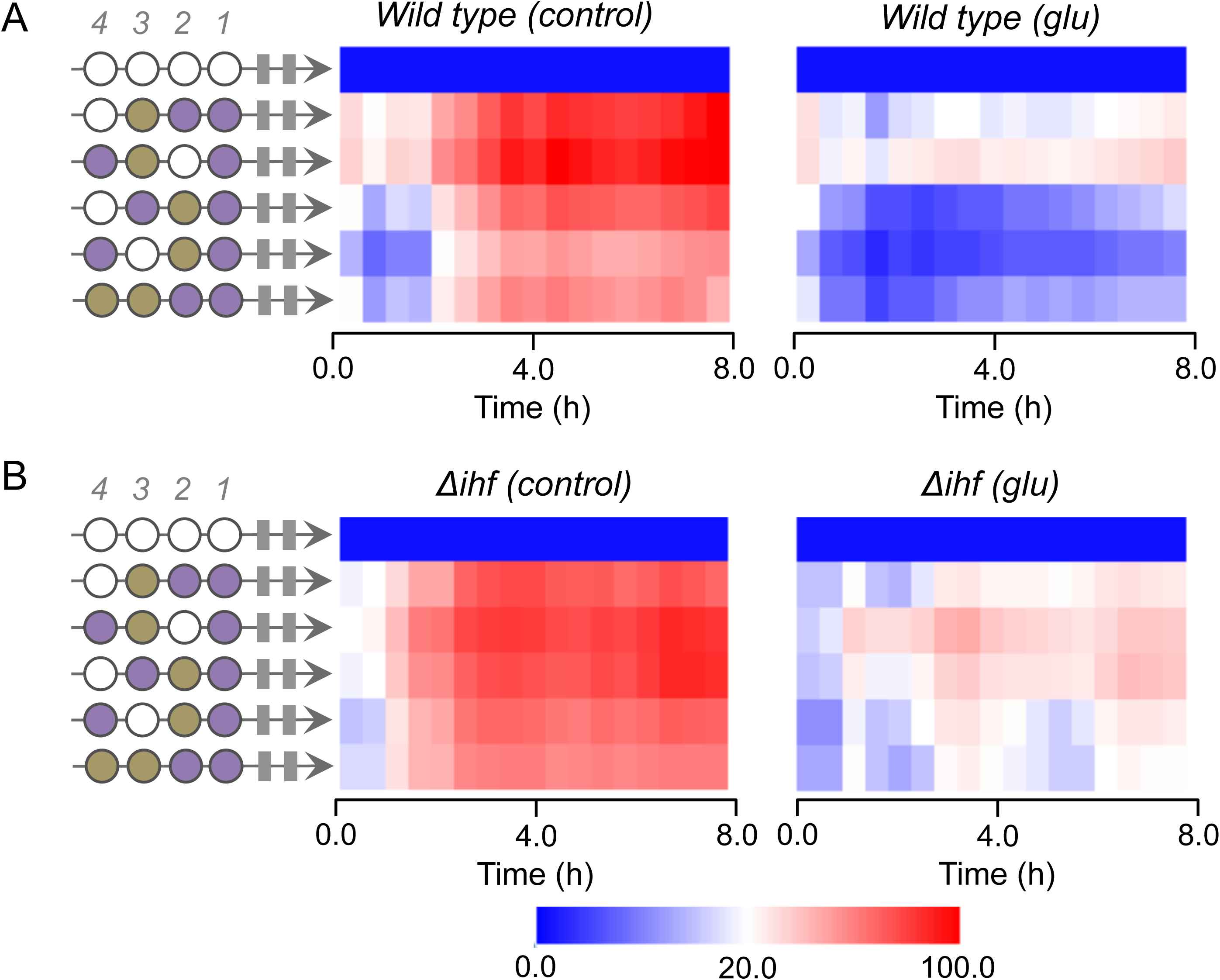
Fine-tuning of CRP-dependent synthetic promoters by IHF sites. Synthetic promoters with the CRP site fixed at position 1 and varying CRP and IHF sites at positions 4, 3, and 2 were assayed in the absence (left) or presence (right) of 0.4% glucose. A) Analysis of promoter activity in the *E. coli* wild type strain. B) Analysis of promoter activity in the *E. coli* Δ*ihf* strain. Relative promoter activity was measured for 8 h, calculated based on the Neutral full promoter, and displayed on an intensity scale from 0.0 to 100.0. Plots were calculated based on the average of three independent experiments.

## Discussion

Regulation of gene expression at the level of RNAP recruitment to target promoters is known to be a combinatorial mechanism where multiple transcriptional factors binding to target *cis*-regulatory elements and their interplay defines the timing and intensity of gene expression. This combinatorial control has been extensively described in bacteria and in single-celled and multicellular eukaryotes, and the so-called *regulatory code* is known to play a major role in the way living organisms develop and interact with the environment (Kinkhabwala & Guet, 2008; Raveh-Sadka et al, 2012; Salgado et al, 2013). However, while classical approaches to understand this code are based on a case-by-case dissection of the *cis*-regulatory elements of particular genes, several studies have now described the systematic investigation of combinatorial promoters through the construction and evaluation of synthetic promoters built from *cis*-regulatory elements. In this sense, Cox III and colleagues constructed a library of synthetic promoters for two local activators (AraC and LuxR) and two local repressors (LacI and TetR) at three different promoter positions (upstream, downstream, or overlapping the core -35/-10 box). From this work, the authors described a number of rules for engineering combinatorial promoters for synthetic biology; for instance, activators were only efficient upstream of the core whereas efficacy of repression was higher at the core and then at the downstream region, with only minor effects at the upstream position (Cox et al, 2007). However, this work only used local TFs, which are limited to a few natural targets and, thus, are not found in naturally complex promoter architectures as global regulators are. Moreover, the work by Cox III only explored a single binding site at the upstream promoter regions, which does not allow the investigation of combinatorial effects generated by *cis*-element arrangements and identities in this region. Therefore, our work addresses a more realistic combinatorial situation by mimicking the manner in which promoters are organized naturally, and indeed, our result of *cis*-element mediated repression of gene expression has not been reported previously. The effect of promoter architecture in gene regulation has also been extensively investigated in single-celled eukaryotes such as yeast, with especial interest in the work of Sharon and co-workers (Sharon et al, 2012). In this study, the authors synthesized and analyzed using a high-throughput approach, thousands of different promoters for several TFs of *Saccharomyces cerevisiae* (Sharon et al, 2012), thus allowing them to investigate the effect of number, position, and affinity of binding sites on gene expression. However, the fundamental difference between transcription initiation in prokaryotes and eukaryotes, due to the sophisticated process of chromatin remodeling required in the latter, makes it impossible to extrapolate the conclusions drawn by Sharon *et al* to a bacterial organism. However, the approach used in this study was analogous to the approach used by Sharon *et al*, since we could inspect the effect of binding site multiplicity, location, and identity.

From the results generated in this work, the most striking was the observation that a single CRP-binding site located immediately upstream of an IHF-binding site could completely abolish transcriptional activity independently of CRP function. This result appeared in several promoter architectures tested here and would indicate that the DNA sequence itself was modulating gene expression. However, introduction of an additional CRP binding site drastically changed this process, resulting in a CRP-activated promoter. It has now been widely demonstrated that DNA can display an allosteric effect on TFs, where the binding of a protein to DNA changes the way this protein interacts with other TFs (Chaires, 2008; Kim et al, 2013; Lefstin & Yamamoto, 1998). Moreover, another type of DNA-based allosteric event has been described where the binding of a protein to DNA can influence the binding of a second protein to an adjacent site independently of protein-protein interaction, and that this influence is transmitted through the DNA molecule (Chaires, 2008; Lefstin & Yamamoto, 1998). In this sense, these processes could explain how two tandem *cis*-elements for CRP that are inactive alone (at positions 3 and 2 in **Fig. 2**) generated a strong CRP-dependent promoter when in association with a single IHF binding site (the latter at position 1, **Fig. 5B**). However, it certainly does not explain how a single CRP *cis*-element displays inhibitory effects in certain promoter architectures (as in many of those presented in **Fig. 4**). Recently, an increasing number of reports have demonstrated that flanking DNA sequences can strongly affect the binding affinity of eukaryotic TFs for identical binding sites (Gordan et al, 2013; Khoueiry et al, 2010), thus explaining why *in vitro* and *in vivo* binding assays do not always correlate. In this process, these flanking sequences generate distortions in the local DNA shape that influences the way the TFs interacts with DNA, by altering the groove width and helical parameters of DNA (Gordan et al, 2013). Though we could not find any report of this process influencing bacterial TFs, our results on synthetic complex promoters suggest that a similar process could influence the activity of bacterial promoters, thus explaining the intrinsic repressive activity of the CRP *cis*-element (independently of the presence of CRP protein) at some positions in promoters containing *cis*-elements for IHF. Our findings could thus be extended to naturally complex promoters and indicate that in those systems, not only would the nature of the TF recruited to the target promoter be imperative for gene expression, but also the *cis*-element itself could have a regulatory role in proximal sites. This evidences an unanticipated intrinsic complexity of natural bacterial promoters that should be considered both for synthetic biology projects as well as to understand the regulatory behavior of natural strains. Taken together, our results highlight the appearance of emergent properties in combinatorial control in bacteria, thus opening new venues for understanding combinatorial regulation in bacterial genes and open new venues that could be investigated in future studies.

## Author Contributions

Conceptualization, R.S.R; Methodology, R.S.R, and L.M.A; Investigation, L.M.O.M., and L.M.A.; Writing – Original Draft, L.M.O.M., and L.M.A.; Writing–Review & Editing, L.M.O.M, and R.S.R.; Funding Acquisition, R.S.R.; Resources, R.S.R.; Supervision, R.S.R.

## Acknowledgments

The authors are thankful to lab members and to Dr. Maria-Eugenia Guazzaroni for critical discussion of this work. This work was supported by the Young Investigator Award of the Sao Paulo State Foundation (FAPESP, award number 2012/22921-8). L.M.O.M. was supported by a FAPESP PhD fellowship (FAPESP, award number 2016/19179-9). The authors declare no conflict of interest.

---

**Table 1.**
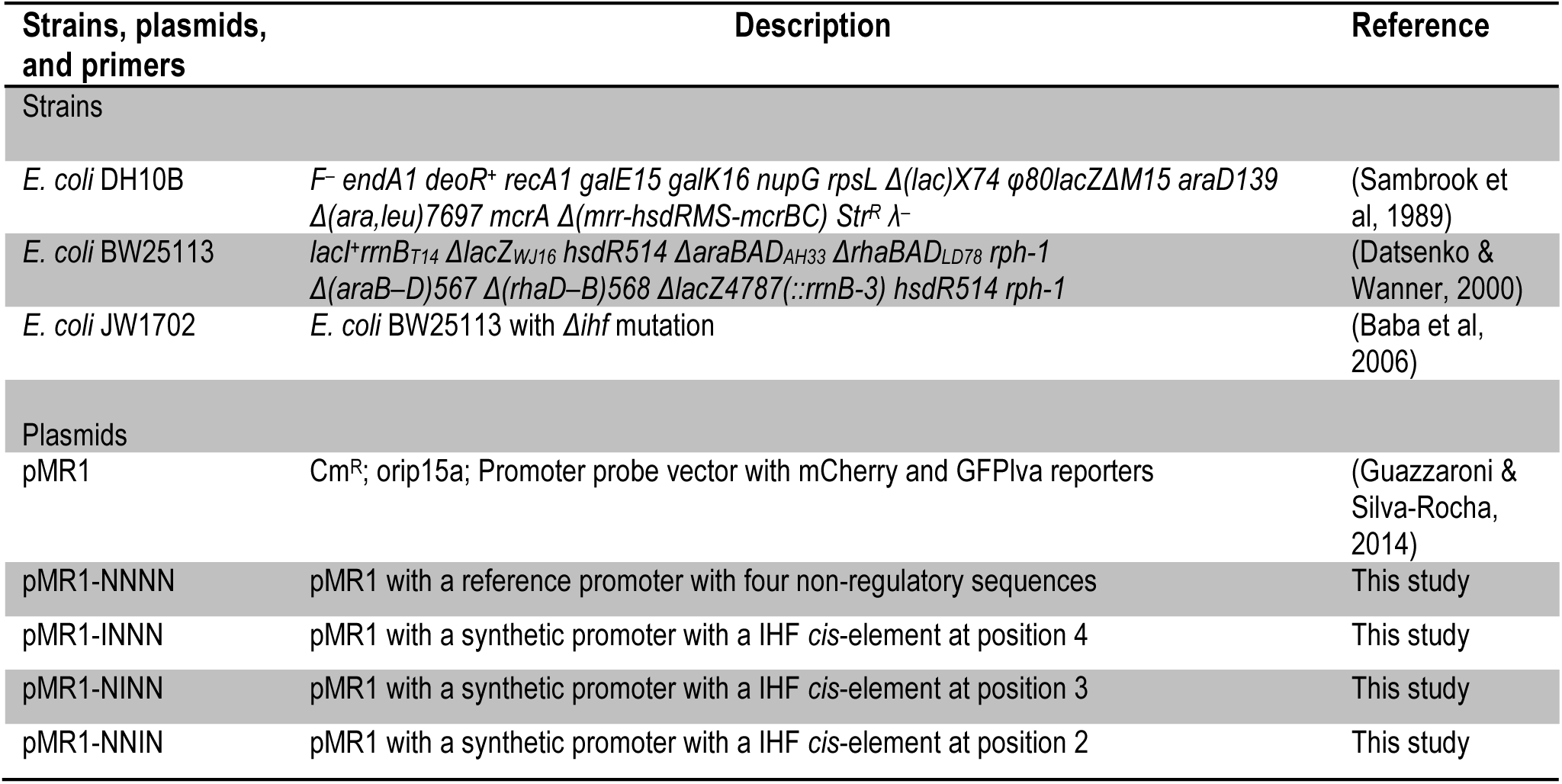
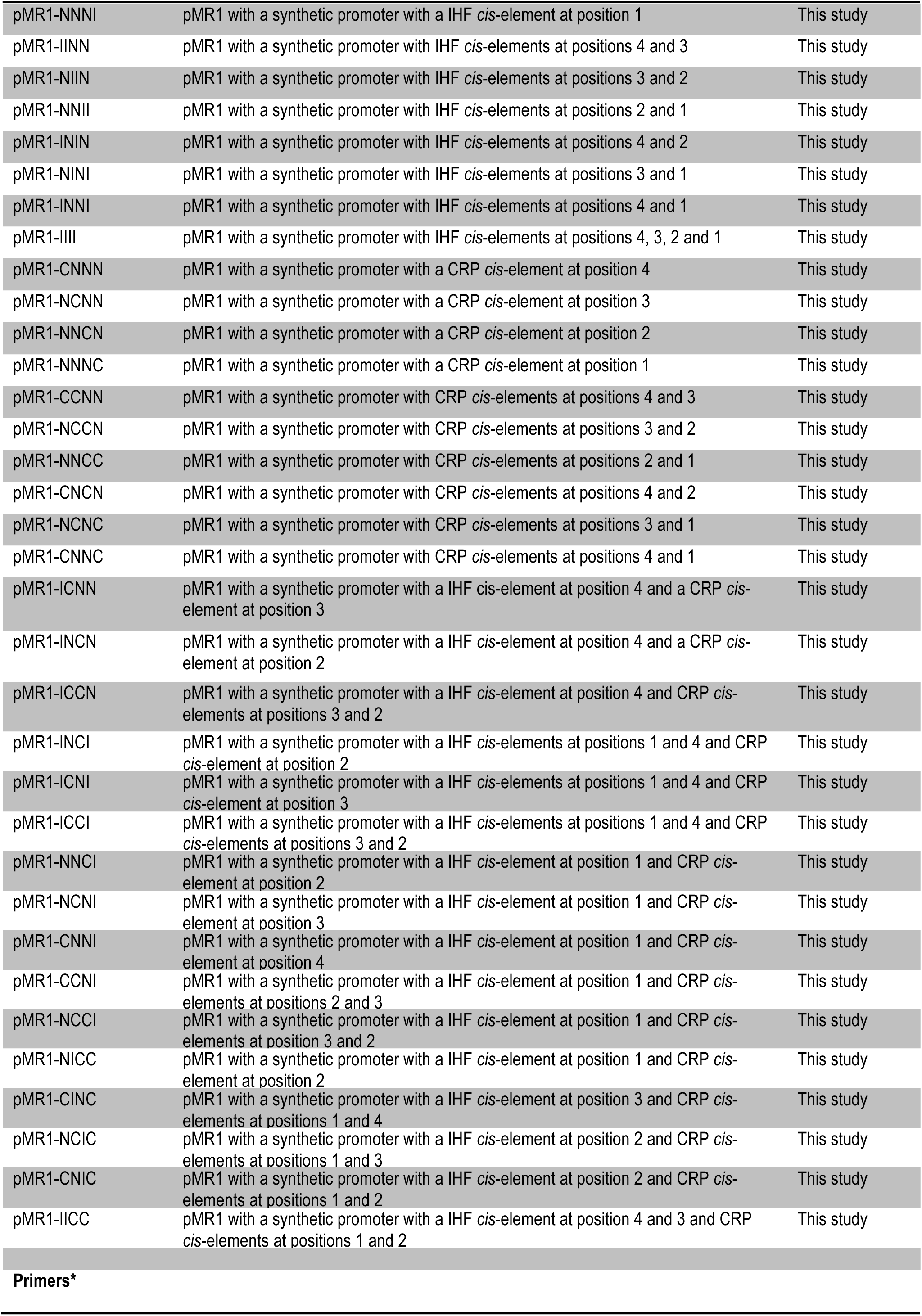
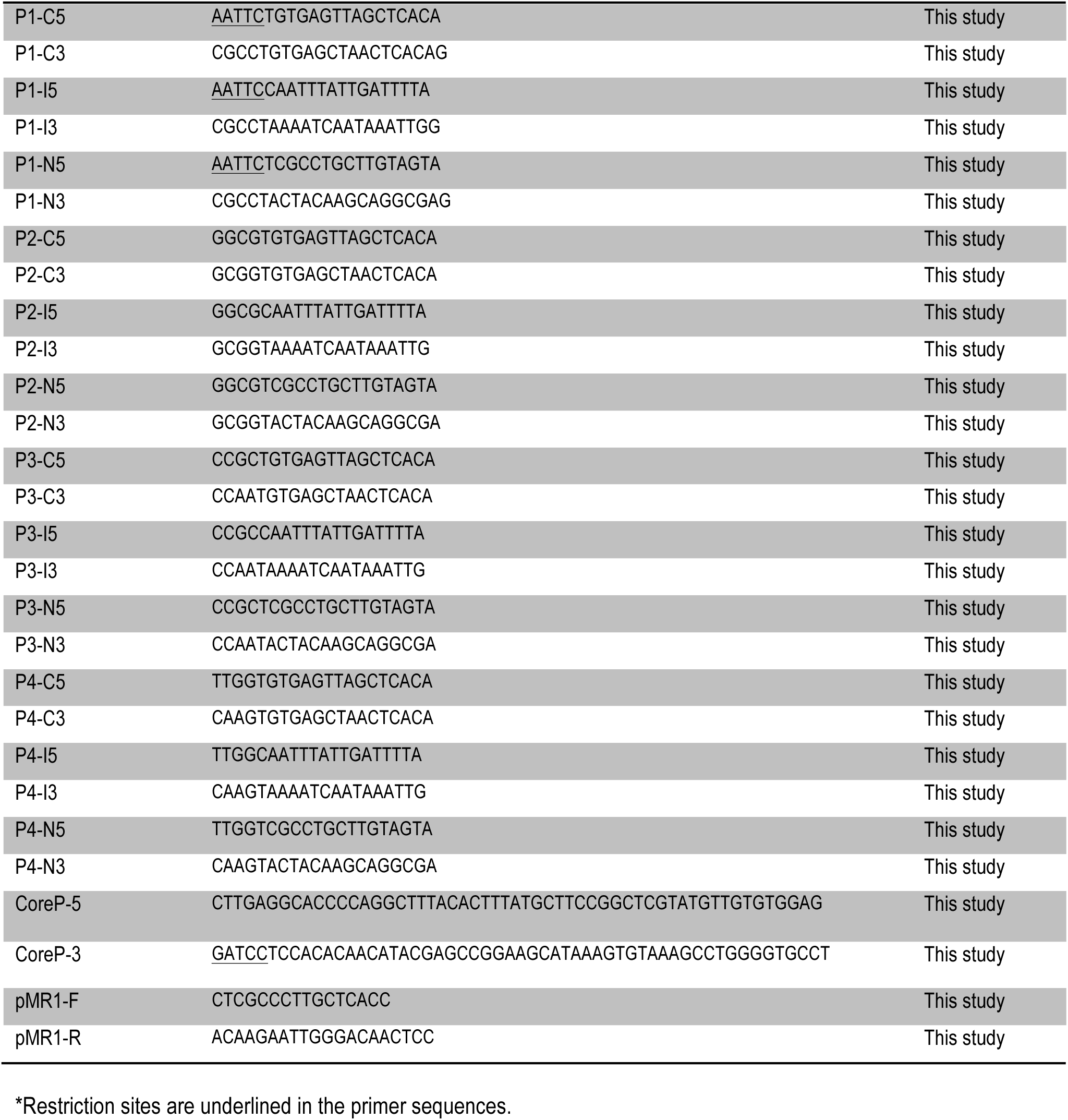
Strains, plasmids and primers used in this study.

## Materials and Methods

### Plasmids, bacterial strains, and growth conditions

The plasmids, bacterial strains, and primers used in this study are listed in Table S1. For cloning procedures, the bacterial strain used was *E. coli* DH5α. *E. coli* BW25113 was used as the wild type strain (WT) whereas *E. coli* JW1702-1 was used as the mutant for IHF transcription factor, and both were obtained from the Keio collection (Baba et al, 2006). *E. coli* strains were grown at 37°C in LB media with chloramphenicol at 34 μg mL^-1^ or in M9 minimal media (6.4 g L^-1^ Na_2_HPO_4_•7H_2_O, 1.5 g L^-1^ KH_2_PO_4_, 0.25 g L^-1^ NaCl, 0.5 g L^-1^ NH_4_Cl) supplemented with chloramphenicol at 17 μg mL^-1^, 2mM MgSO_4_, 0.1mM casamino acids, and 1% glycerol as the sole carbon source. Where indicated, CRP response was depleted by using 0.4% of glucose.

### Design of the minimal promoter scaffold and ligation reactions

Promoters were constructed by ligation of 5′ end phosphorylated oligonucleotides (Cox et al, 2007; Kinkhabwala & Guet, 2008) acquired from Sigma Aldrich (**Table S1**). All single strand nucleotides were designed to carry a discrete 16 bp sequence^8^ containing a CRP binding site (C), IHF binding site (I), one Neutral (N) motif with no transcription factor binding (**Fig. 1B**), and a core promoter based on the *lac* promoter (**Table S1**), which is a weak promoter and therefore requires activation. All these oligonucleotides were designed to carry three base pair overhangs corresponding to their corrected insertion region on the promoter (**Fig. 1A**). The upper and lower strand corresponding to each position were mixed at equimolar concentrations and annealed by heating at 95°C followed by gradual cooling to room temperature. External overhangs of the fourth *cis*-element position and the core promoters reassembled on the EcoRI and BamHI digested sites, allowing ligation to a previously digested EcoRI/BamHI pMR1 (Guazzaroni & Silva-Rocha, 2014) plasmid. All five fragments (four *cis*-elements positions plus core promoter) were mixed at equimolar concentrations in a pool with the final concentration of 5′ phosphate termini fixed at 15 μM. For the ligase reaction, 1 μL of the pooled fragments was added to 50 ng EcoRI/BamHI pMR1 digested plasmid in presence of ligase buffer and ligase enzyme to a final volume of 10 μL. After one hour at 16°C, the ligase reaction was inactivated for 15 min at 65°C and one aliquot of 2 μL was then electroporated into 50 μL of *E. coli* DH10B competent cells. After one hour of regenerating in 1 mL LB media, the total volume was plated onto LB solid dishes supplemented with chloramphenicol at 34 μg mL^-1^. Clones were confirmed by colony PCR using primers pMR1-F and pMR1-R (**Table S1**) using the pMR1 empty plasmid PCR reaction as a further length reference upon agarose gel electrophoresis. Clones with the potential correct length were submitted to Sanger DNA sequencing for confirming the correct promoter assembly.

### GFP fluorescence assay and data processing

To measure promoter activity, the library of 38 promoters was analyzed in different genetic backgrounds and conditions. For each experiment, a plasmid harboring the promoter of interest was used to transform *E. coli wild type or E. coli* Δ*ihf* mutant cells. Freshly plated single colonies were grown overnight in LB media, centrifuged, and resuspended in fresh M9 media. The culture (10 μL) was then assayed in 96-well microplates in biological triplicates with 170 μL of M9 media or M9 media supplemented with 0.4% glucose whenever required. Cell growth and GFP fluorescence were quantified using a Victor X3 plate reader (Perkin Elmer). Promoter response was calculated as arbitrary units by dividing the fluorescence levels by the optical density at 600 nm (reported as GFP/OD600) after background correction. The same strain harboring the pMR1 empty plasmid was used as the threshold background signal during calculations. Fluorescence and absorbance measurements were taken at 30 min intervals over 8 h. Technical triplicates and biological triplicates were included in all experiments. Raw data were processed using *ad hoc* R script (https://www.r-project.org/) and plots were constructed using R or MeV (www.tm4.org/mev.html).

